# Improvements in the Sequencing and Assembly of Plant Genomes

**DOI:** 10.1101/2021.01.22.427724

**Authors:** Priyanka Sharma, Othman Al-Dossary, Bader Alsubaie, Ibrahim Al-Mssallem, Onkar Nath, Neena Mitter, Gabriel Rodrigues Alves Margarido, Bruce Topp, Valentine Murigneux, Ardy Kharabian Masouleh, Agnelo Furtado, Robert J Henry

## Abstract

**Background:** Advances in DNA sequencing have reduced the difficulty of sequencing and assembling plant genomes. A range of methods for long read sequencing and assembly have been recently compared and we now extend the earlier study and report a comparison with more recent methods.

**Results:** Updated Oxford Nanopore Technology software supported improved assemblies. The use of more accurate sequences produced by repeated sequencing of the same molecule (PacBio HiFi) resulted in much less fragmented assembly of sequencing reads. The use of more data to give increased genome coverage resulted in longer contigs (higher N50) but reduced the total length of the assemblies and improved genome completeness (BUSCO). The original model species, *Macadamia jansenii*, a basal eudicot, was also compared with the 3 other *Macadamia* species and with avocado (*Persea americana*), a magnoliid, and jojoba (*Simmondsia chinensis*) a core eudicot. In these phylogenetically diverse angiosperms, increasing sequence data volumes also caused a highly linear increase in contig size, decreased assembly length and further improved already high completeness. Differences in genome size and sequence complexity apparently influenced the success of assembly from these different species.

**Conclusions:** Advances in long read sequencing technology have continued to significantly improve the results of sequencing and assembly of plant genomes. However, results were consistently improved by greater genome coverage (using an increased number of reads) with the amount needed to achieve a particular level of assembly being species dependant.

## Background

Recent advances in DNA sequencing technology have facilitated the sequencing and assembly of plant genomes with a rapid growth in reports of high quality chromosome level assemblies [1]. A basal eudicot, *Macadamia jansenii*, was used to compare the range of long read sequencing and assembly technologies available in 2019[2]. The Pac Bio (Sequel), Oxford Nanopore Technology (PromethION) and BGI (single-tube Long Fragment Read) platforms were applied to the analysis of the same sample. Assembly tools were evaluated for these data sets and the contribution of short reads to improving assemblies assessed [2]. Technology improvements had delivered ongoing increases in the length and quality of sequence reads delivered by these platforms. Since the original study, significant further advances have been made with the use of repeated sequencing of the same molecule to greatly increase sequence accuracy for long reads. This technology allows the generation of long reads (10-25kb) with greater than 99.5% accuracy [3]. Comparison of long read technologies demonstrates the pros and cons of different platforms in relation to contiguity, accuracy of sequence and data analysis time [4]. We now update the earlier study to demonstrate the impact of these improvements on genome assemblies. Factors such as the volume of data (bp) used in the assembly were explored for the *Macadamia* genome, related species and other diverse species with similar sized genomes.

### Long read versus HiFi assemblies

Comparison of assemblies based upon long reads [5] and circular consensus sequence (CCS) reads from HiFi [3], showed the greater accuracy of the CCS reads resulted in greatly improved assemblies for the *Macadamia jansenii* genome (Table 1).

**Table 1.** Improvement in long read sequencing (Pac Bio) for *Macadamia jansenii* when using higher accuracy sequencing.

|  | Long reads*[2] | HiFi |
| --- | --- | --- |
| Total data | 65.2Gb | 28Gb |
| Contig N50 | 1.57Mb | 4.49Mb |
| Assembly length | 758Mb | 738Mb |
| Number of contigs | 762 | 284 |
| BUSCO | 97% | 98% |

\*phased Falcon assembly

The assembly with the high quality HiFi reads was less fragmented with slightly reduced total genome length and improved completeness (BUSCO). The use of around 20Gb of high quality (HiFi) data gave N50 values of 4 Mb and resulted in assemblies with fewer than 300 contigs required to cover the genome. This represents a significant advance over the assemblies possible when this sample was used previously to compare different long read platforms and assembly tools, many of which required long computing times to assemble contigs [2]. The high quality IPA assembly had a run time of 20 h with 120 Gb peak memory on the FlashLite computer cluster. This analysis requirement compares favourably with the results for a large number of earlier assembly tools reported for the same sample[2], but provides a much higher quality assembly. Assembly of the HiFi data with other recent tools was also compared. Flye resulted in a highly fragment genome of 993 Mb with an N50 of 459 kb, while Hifiasm produced a genome of 827 Mb composed of 779 contigs but with an N50 of 46.1 Mb and a L75 of 14.

### Results for other *Macadamia* species

*Macadami jansenii* is an endangered species and is one of four species in the *Macadamia* genus. Sequences of all four species were obtained using the same HiFi techniques and all gave similar high quality outcomes when assembled (Table 2).

**Table 2.** Comparison of assemblies of *Macadamia* species*

|  | *M. janseni* | *M. integrifolia* | *M. tetraphylla* | *M. ternifolia* |
| --- | --- | --- | --- | --- |
| Contig N50 | 4.5Mb | 5.3Mb | 10.0Mb | 6.4Mb |
| Longest Contig | 16.6Mb | 26.4Mb | 32.1Mb | 21.2Mb |
| Assembly Length | 738Mb | 742Mb | 707Mb | 716Mb |
| Number of contigs | 284 | 249 | 153 | 211 |
| BUSCO | 98% | 98% | 97% | 98% |

\*Primary assemblies shown. For details of associate assemblies see supplementary Table 1.

### Results for other diverse plant species

Methods for sequencing plant genomes need to be applied to genomes with a wide range of sizes and complexities. Macadamia is a basal eudicot. Other diverse flowering plant genomes were sequenced to determine how widely applicable the results of this study would be in plant genome sequencing. Jojoba (*Simmondsia chinensis*), a core eudicot from the Caryopyllales, and avocado, a magnoliid, were compared. The three diverse genomes were all similar in size (700-1000Mb). Many important plant genomes are in or near this size range [6]. *M. jansenii* is an endangered species with relatively low heterozygosity, avocado has much greater heterozygosity [7] and jojoba has been reported to be a tetraploid[8]. Heterozygosity and polyploidy both complicate assembly [9,10]. The quality of the assemblies was more contiguous (fewer contigs required to cover the genome) or similar (avocado) with less data in each of these cases when HiFi reads were used instead of the earlier continuous long reads (Table 3). The macadamia and jojoba genomes gave N50 values that were larger when using the HiFi (CCS) reads than with long reads (CLR). However, the N50 for the slightly larger genome of avocado was greater when using the long reads compared to that obtained with the HiFi reads. This suggest that the larger genome may have longer repeat regions that limit contig assembly in some parts of the genome with HiFi reads.

**Table 3.** Long read versus HiFi sequencing of other diverse species

|  | Long reads |  | HiFi |  |
| --- | --- | --- | --- | --- |
|  | Jojoba | Avocado | Jojoba | Avocado |
| Total data | 152Gb | 159Gb | 41.4Gb | 44Gb |
| Contig N50 | 4.73Mb | 6.7Mb | 4.89Mb | 4.3Mb |
| Assembly length | 1260Mb | 787Mb | 780Mb | 749Mb |
| Number of contigs | 762 | 308 | 284 | 298 |
| BUSCO | 99% | 99% | 98% | 98% |

### Impact of the sequencing coverage on the assemblies

The length of the contigs assembled (Figure1) was directly related to the volume of sequence data used. Analysis of four related Macadamia species gave a similar linear relationship between data volume and contig N50 for input of between 10 and 40 times genome coverage. The size of the contigs assembled showed a similar dependence upon the amount of sequence data (genome coverage) across species with the slope of the relationship varying for different species (Figure 2). The macadamia genomes could be assembled with lower coverage. This may be a function of genome size with their smaller genomes requiring less coverage to achieve a given N50. The larger genomes may contain a higher proportion of repetitive sequences that are difficult to assemble.

**Figure 1.**
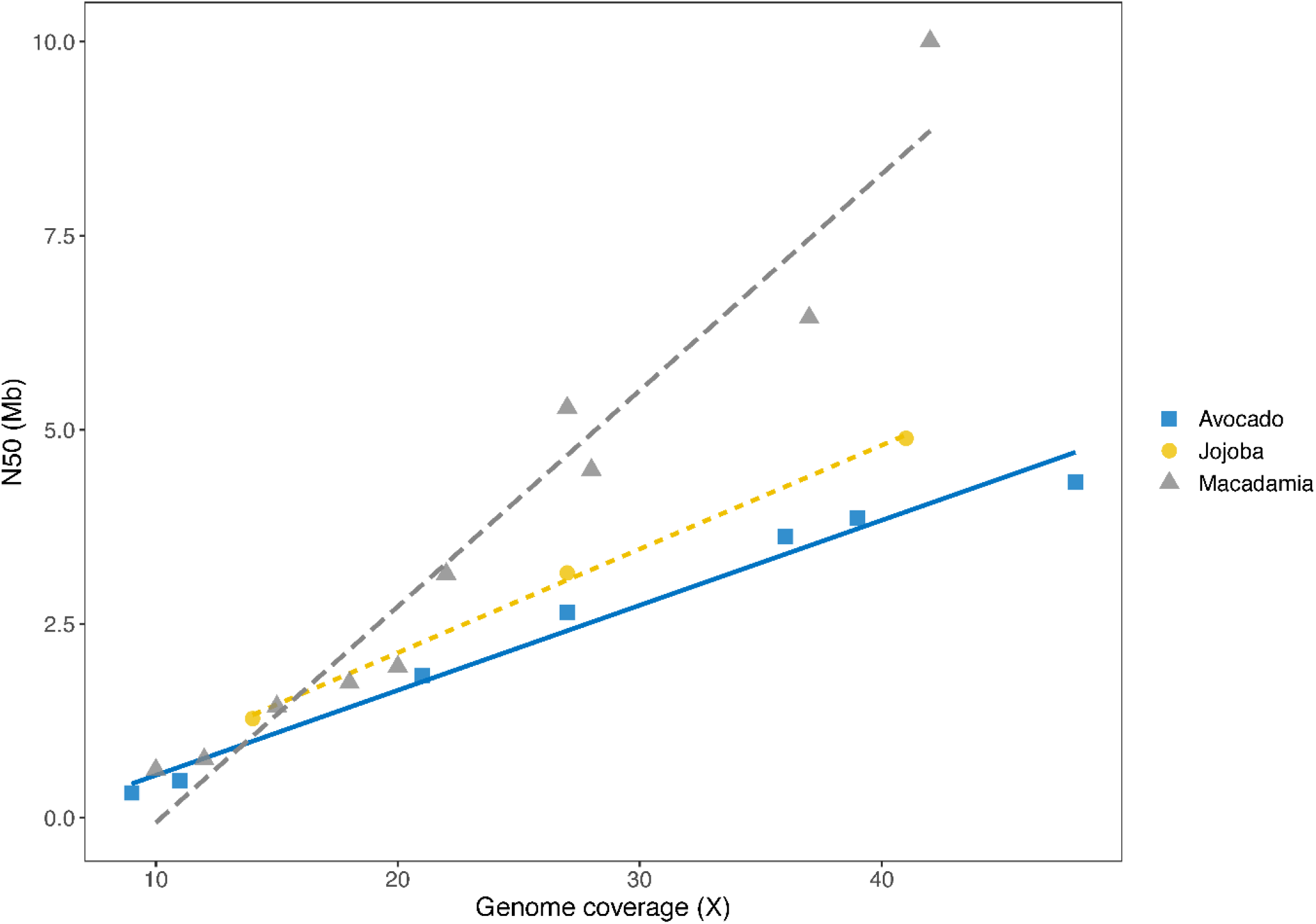
Influence of data volume on assembly for Macadamia species N50 of contigs is plotted against the genome coverage. Genome sizes used to calculate coverage were; *M*.*integrifolia*, 895Mb [12]; *M janseni*, 780Mb [2]; *M. tetraphylla*, 758Mb [23] and *M. ternifolia*, 758Mb (not known but assumed the same as *M. tetraphylla* due to similar assembly size).

**Figure 2.**
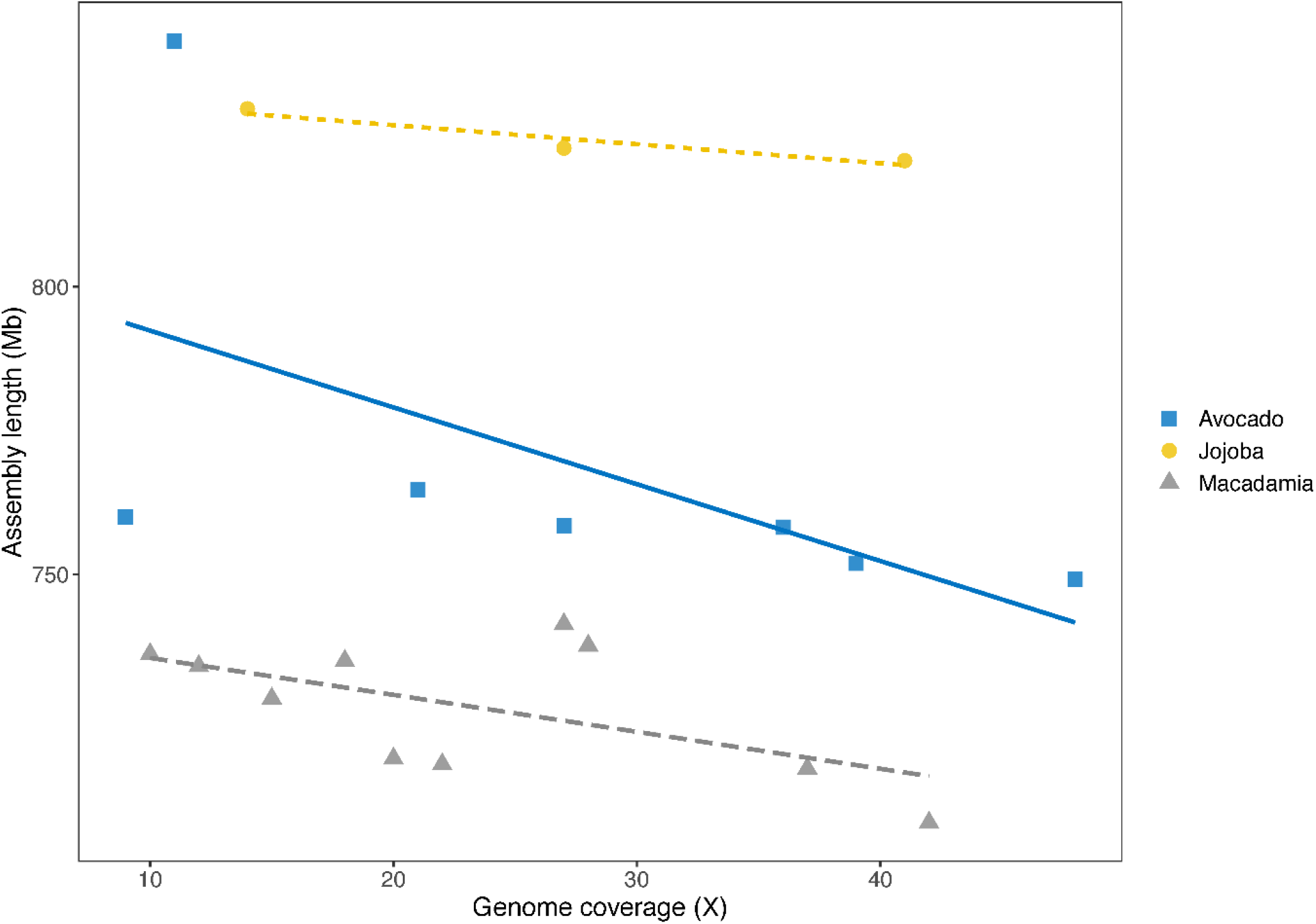
Influence of data volume on assembly for diverse species. N50 of contigs is plotted against the genome coverage. Genome sizes used to calculate coverage were, jojoba, 1003Mb[14]; avocado, 920Mb [24] and as in figure 1 for *Macadamia* species

Assemblies based upon more data were slightly shorter in total length (Figure 3). This reduction was probably due to removal of duplicated end sequences as contigs were joined. The high quality of these assemblies was confirmed by BUSCO values of more than 95%. Genome completeness was high in all cases but increased slightly when more data was used in the assembly (Figure 4).

**Figure 3.**
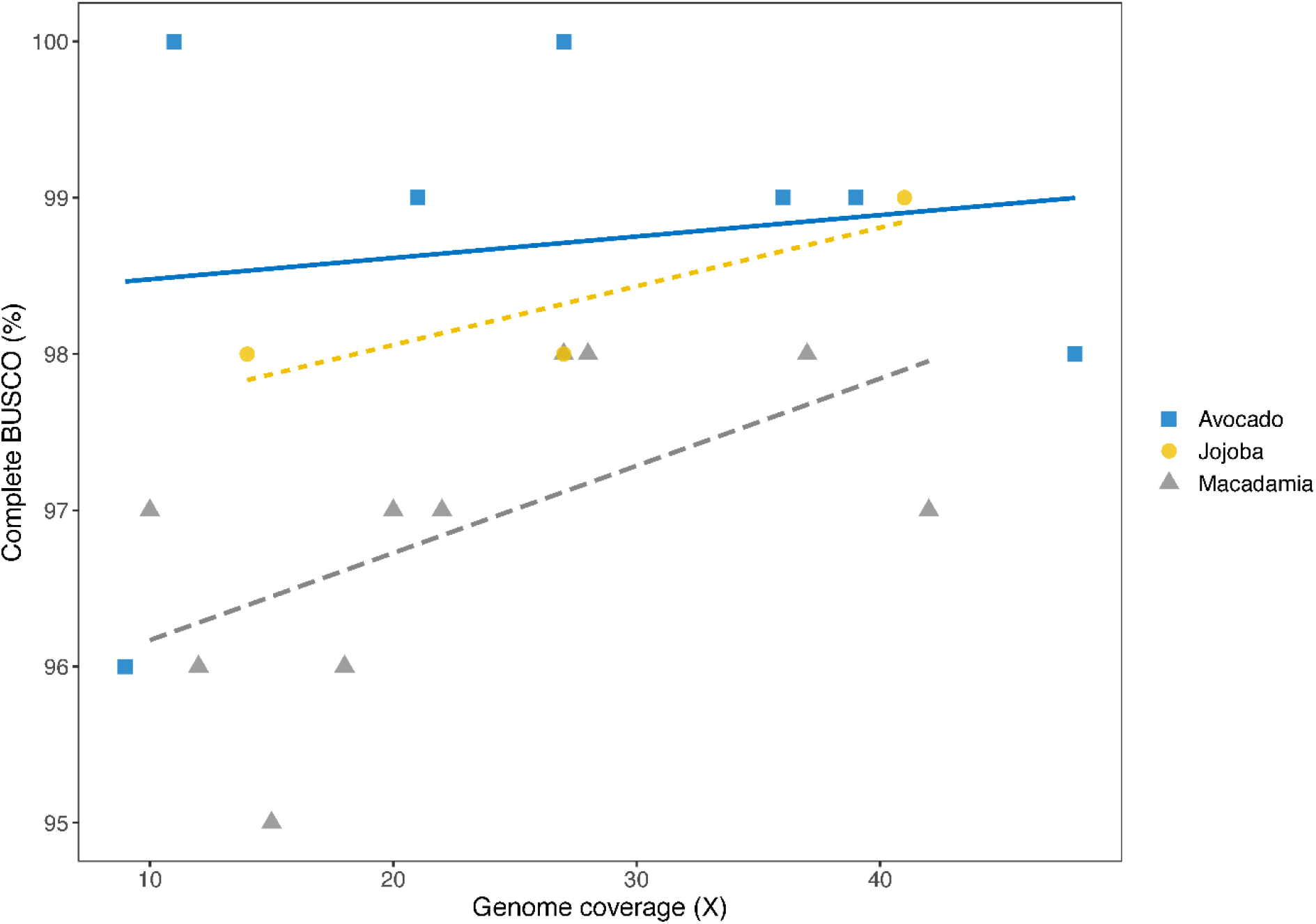
Decrease in length of total assembly as more genome coverage is used in the assembly

**Figure 4.**
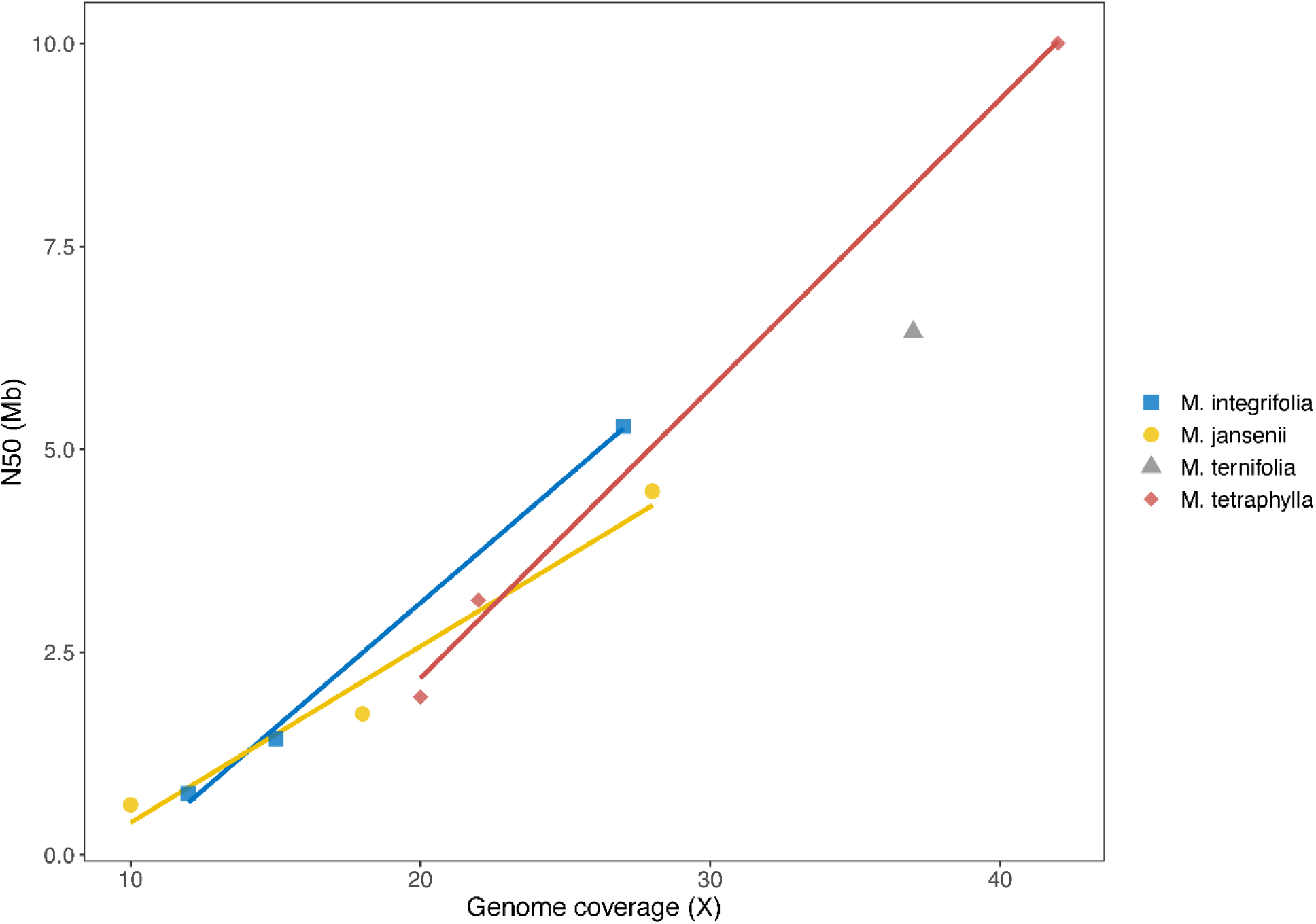
Improvement in genome completeness (BUSCO%) with genome coverage

These results were confirmed when applying these methods to sequencing the other phylogenetically diverse plant genomes with slightly larger genomes with greater genome complexity. In each case N50 and completeness increased with data volume and genome size declined.

### Impact of the read length on the assemblies

The length of sequence reads was also expected to influence the assembly. Examination of size distribution of the 6 species showed that the length of the sequences varied slightly within the expected range around 15kb for HiFi data. The minor differences in mean read length and numbers of longer reads did not explain the differences in the size of the contigs assembled (Supplementary Figure 1). This suggest that the different amounts of sequence data required to drive assembly to a particular level may be associated more with the complexity of the sequence. The close relationship between sequence volume and N50 for the four *Macadamia* species may reflect the similar sequence complexity of the species in this group. The jojoba and avocado required more sequence data to reach the same level of assembly. The slightly larger genome size of these two species may be enough to explain this difference due to the likely higher proportion of repetitive sequence in the somewhat larger genomes.

### Oxford Nanopore Technologies updates

Oxford Nanopore Technologies (ONT) regularly releases updated basecalling software to convert the raw electrical signal into sequence data. We repeated the basecalling of the ONT raw data of *M. jansenii* using different versions of the Guppy basecaller released in March 2019 (v2.3.7), April 2019 (v3.0.3) and June 2020 (v4.0.11). The assembly quality improved as shown by an increase in the assembly contiguity and in the number of complete BUSCOs before any polishing (Table 4). The assembly size decreased from 817 Mb to 798 Mb. Two versions of the Flye assembler were applied to the same basecalled sequence dataset, which resulted in a significant increase in genome contiguity and completeness as well as a reduced genome assembly size.

**Table 4.**
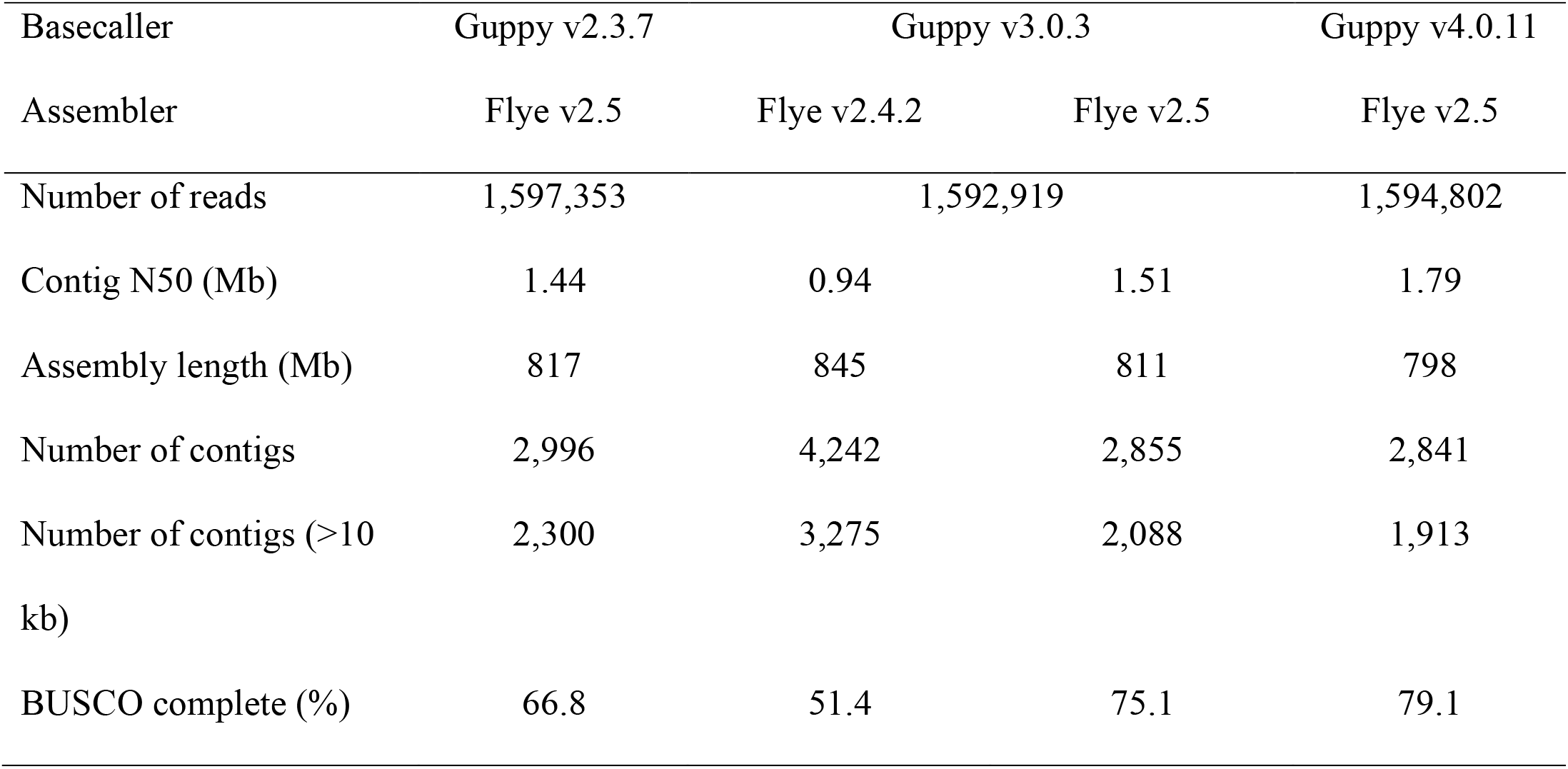
ONT genome assembly statistics of *M. jansenii* using the Flye assembler, the pass reads and different Guppy basecallers versions

### Conclusions

These assemblies represent significant advances over the highly fragmented genomes previously reported for these species [11-14]. Advances in long read sequencing using different platforms provide improving options for plant genome sequencing and assembly [15]. A recent comparison of these methods applied to rice genome sequencing showed strengths and weaknesses of both with greater sequence accuracy in the Pac Bio assemblies and more contiguity in the ONT assemblies [4]. The resulting genome sequences can be evaluated for the best combination of sequence and assembly accuracy [16]. The results presented here show that contig size can be increased by adding more sequence reads to achieve a linear increase in N50. This extra data will result in slightly shorter total assembly lengths and improved completeness of the genomes. The improved methods when combined with higher level assembly tools[17] will support routine, rapid and efficient generation of highly accurate chromosome level genome sequences of plant species[18].

## Methods

### DNA extraction

All local, national and international guidelines and legislation was observed in obtaining samples for this study. *Macadamia jansenii* DNA was prepared as described earlier[19]. Three other Macadamia species (*M. tetraphylla, M. ternifolia* and *M. integrifolia*) and Jojoba (*Simmondsia chinensis*) were also extracted using this method with minor modifications where phenol was excluded from the extraction method[20]. Avocado (*Persea americana*) DNA was isolated by a modified CTAB (cetyl-trimethyl ammonium bromide) DNA extraction protocol [21,22]. Leaf tissue (0.2 g) was ground and added to 15 ml of 2% CTAB buffer, pH 8.0 followed by 15 min incubation at 65 °C. The supernatant after centrifugation at 10 g for 15 min was treated with RNAse A (10ng/µl) and incubated at 37°C for 30 min. Chloroform: Isoamyl alcohol (24:1) washes were performed followed by precipitation with isopropanol and 70% ethanol washes. The DNA was resuspended in ultrapure DNAse and RNAse free water for sequencing.

### DNA sequencing and assembly

Long read sequencing was as previously described[19]. Long reads (CLR) were assembled using Falcon [2] for *M. jansenii* and Canu v 2.0 for the other genomes. HiFi gDNA libraries were prepared using sheared genomic DNA (∼15-20 kb) was sequenced on a PacBio Sequel II (software/chemistry v9.0.0) following diffusion loading. Sequence data was processed to generate CCS reads using the default settings of the CCS application (v4.2.0) in SMRT Link (v9.0.0); minimum parameters for passes (3), accuracy (0.99), CCS read length (10) and maximum CCS read length (50000). CCS reads were assembled using the Improved Phased Assembly (IPA) method (PacBio). The IPA assembly tool (https://github.com/PacificBiosciences/pbbioconda/wiki/Improved-Phased-Assembler) uses the HiFi sequencing reads (high-quality consensus reads) and generates phased assembly. This produces a primary contig folder, including the main assembly and an associated contig, containing haplotigs and duplications. For all assemblies, 24 CPUs and 120Gb of memory was employed.

### Assessment of completeness

The completeness of genome assemblies was evaluated using benchmarking universal single-copy orthologues (BUSCO) analysis (v4.1.2), using genome mode and lineage Eukaryota_odb10 dataset.

## Availability of data

Sequence data from Pac Bio (Sequel), Oxford Nanopore Technology (PromethION) and BGI (single-tube Long Fragment Read) analysis of *M. jansenii* was described by Murigneux et al [2]. BGI, PacBio, ONT, and Illumina sequencing data generated in that study were deposited in the SRA under BioProject PRJNA609013and BioSample SAMN14217788. Accession numbers are as follows: BGI (SRR11191908), PacBio (SRR11191909), ONT PromethION (SRR11191910), ONT MinION (SRR11191911), and Illumina (SRR11191912). Assemblies and other supporting data are available from the *GigaScience* GigaDB repository [25]. Pac Bio HiFi reads described in this paper are deposited as CCS reads under NCBI BioProject ID Macadamia : PRJNA694456; Avocado: PRJNA694184 and Jojoba: PRJNA694450

## Additional files

Figure S1:Size distribution of reads sequenced

## Author’s contributions

Contributions of authors were as follows. RH, AF, VM, AM, IA Conceptualization; PS, OA, BA, ON, VM, AF Data curation; PS, OA, BA,ON, GM, VM, AM, AF Formal Analysis; RH, AF, IA, AM Funding acquisition; PS, OA, BA, ON, NM, VM, AM, AF,RH Investigation; IA, BT, Resources; IA, BT, NM, RH, AM, AF Supervision; RH, PS, ON, AM Writing – original draft; All authors Writing – review & editing

## Acknowledgements

The project was supported by the University of Queensland Research Computing Centre (RCC) and the University of Queensland Genome Innovation Hub.

---

**Figure S1:**
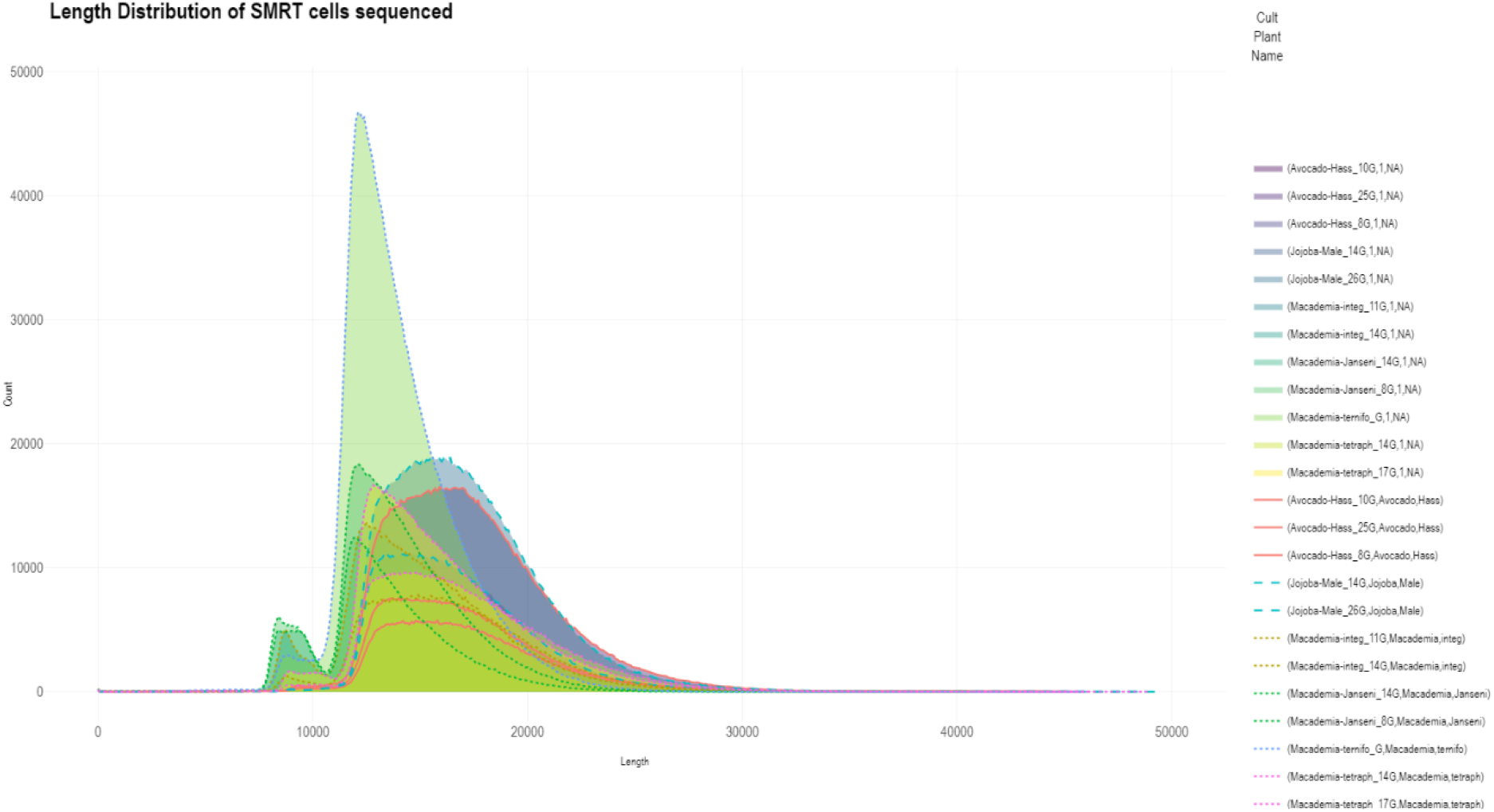
Size distribution of reads sequenced

**Supplementary Table 1.**
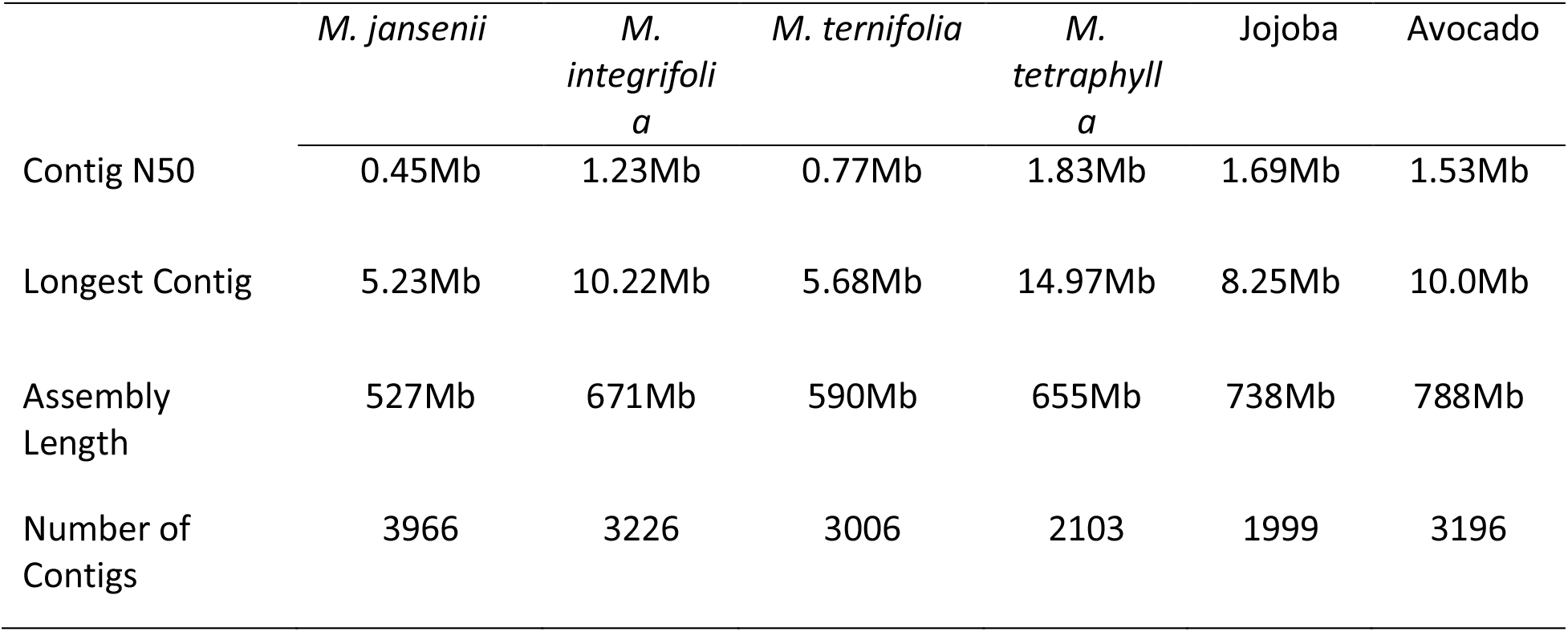
Data for associated contigs in IPA assemblies

